# A mean-field approach to the dynamics of networks of complex neurons, from nonlinear Integrate-and-Fire to Hodgkin-Huxley models

**DOI:** 10.1101/870345

**Authors:** M. Carlu, O. Chehab, L. Dalla Porta, D. Depannemaecker, C. Héricé, M. Jedynak, E. Köksal Ersöz, P. Muratore, S. Souihel, C. Capone, Y. Zerlaut, A. Destexhe, M. di Volo

**Affiliations:** Department of Integrative and Computational Neuroscience, Paris-Saclay Institute of Neuroscience, Centre National de la Recherche Scientifique, 91198 Gif sur Yvette, France; Ecole Normale Superieure Paris-Saclay, France; Institut d’Investigacions Biomèdiques August Pi i Sunyer, Barcelona, Spain; Strathclyde Institute of Pharmacy and Biomedical Sciences, Glasgow, Scotland, UK; Univ. Grenoble Alpes, Inserm, U1216, Grenoble Institut Neurosciences, GIN, 38000 Grenoble, France; Inserm, U1099, 35042 Rennes, France; MathNeuro Team, Inria Sophia Antipolis Méditerranée, 06902 Sophia Antipolis, France; Physics Department, Sapienza University, Rome, Italy; Université Côte d’Azur, Inria Sophia Antipolis Méditerranée, France; Istituto Nazionale di Fisica Nucleare, sezione di Roma, Italy; Laboratoire de Physique Théorique et Modelisation, Université de Cergy-Pontoise, 95302 Cergy-Pontoise cedex, France

**Author notes:** these authors contributed equally (M. Carlu, O. Chehab, L. Dalla Porta, D. Depannemaecker, C. Héricé, M. Jedynak, E. Köksal Ersöz, P. Muratore, S. Souihel).

## Abstract

We present a mean-field formalism able to predict the collective dynamics of large networks of conductance-based interacting spiking neurons. We apply this formalism to several neuronal models, from the simplest Adaptive Exponential Integrate-and-Fire model to the more complex Hodgkin-Huxley and Morris-Lecar models. We show that the resulting mean-field models are capable of predicting the correct spontaneous activity of both excitatory and inhibitory neurons in asynchronous irregular regimes, typical of cortical dynamics. Moreover, it is possible to quantitatively predict the populations response to external stimuli in the form of external spike trains. This mean-field formalism therefore provides a paradigm to bridge the scale between population dynamics and the microscopic complexity of the individual cells physiology.

**NEW & NOTEWORTHY:** Population models are a powerful mathematical tool to study the dynamics of neuronal networks and to simulate the brain at macroscopic scales. We present a mean-field model capable of quantitatively predicting the temporal dynamics of a network of complex spiking neuronal models, from Integrate-and-Fire to Hodgkin-Huxley, thus linking population models to neurons electrophysiology. This opens a perspective on generating biologically realistic mean-field models from electrophysiological recordings.

## 1 Introduction

Brain dynamics can be investigated at different scales, from the microscopic cellular scale, describing the voltage dynamics of neurons and synapses (Markram et al., 2015) to the mesoscopic scale, characterizing the dynamics of whole populations of neurons (Wilson and Cowan, 1972), up to the scale of the whole brain where several populations connect together (Bassett et al., 2018; Deco et al., 2015; Sanz Leon et al., 2013).

In their pioneering work (Wilson and Cowan, 1972), Wilson and Cowan describe the dynamics of a population of neurons through a well-known differential equation where the input-output gain function is described by a sigmoid function. This approach inspired a long lasting research in neuroscience where population models, usually called *rate models*, permit a qualitative insight into the dynamics of population of neurons (di Santo et al., 2018; Hopfield, 1984; Sompolinsky et al., 1988; Sussillo and Abbott, 2009).

Moreover, a large effort has been made in order to derive population descriptions from the specificity of the network model under consideration. This bottom-up approach permits to obtain a dimensionally reduced *mean field* description of the network population dynamics in different regimes (Amit and Brunel, 1997; Brunel and Hakim, 1999; Capone et al., 2019b; di Volo et al., 2014; El Boustani and Destexhe, 2009; Montbrió et al., 2015; Ohira and Cowan, 1993; Renart et al., 2004; Schwalger et al., 2017; Tort-Colet et al., 2019; Tsodyks and Sejnowski, 1995; Van Vreeswijk and Sompolinsky, 1996; Vreeswijk and Sompolinsky, 1998). On one hand *mean field* models permit a simpler, reduced picture of the dynamics of a population of neurons, thus allowing to unveil mechanisms determining specific observed phenomena (di Volo and Torcini, 2018; Jercog et al., 2017; Reig and Sanchez-Vives, 2007). On the other hand, they enable a direct comparison with imaging studies where the spatial resolution implies that the recorded field represents the average over a large population of neurons (i.e. a mean-field) (Capone et al., 2017; Chemla et al., 2019).

During awake states, cortical dynamics generally show asynchronous spiking activity, where individual neurons are characterized by an irregular (typically Poissonian) firing pattern (Burns and Webb, 1976; Dehghani et al., 2016; Softky and Koch, 1993). In this dynamical regime, so-called Asynchronous Irregular (AI), the correlation of the network activity decays relatively quickly in time, making it possible to develop a Markovian formalism in order to obtain mean-field equations. The application of such a theory to binary neurons led to the derivation of dynamical equations for population rates (Ginzburg and Sompolinsky, 1994; Ohira and Cowan, 1993). More recently, such a theory has been extended to spiking neurons, permitting to obtain differential equations for neurons average activity and for higher-order moments (Buice et al., 2010; Dahmen et al., 2016; El Boustani and Destexhe, 2009). In the first order, these equations are formally the same as the *rate models*, like the Wilson-Cowan approach, although the function linking input-output properties of populations of neurons, namely the transfer function, is more complex than a sigmoid. Indeed, in this formalism, it encompasses internal properties of the neuronal models, together with the type of synaptic interactions under consideration, to yield a population scale description. In general, such function cannot be expressed in a closed form for complex neurons, especially if some realistic ingredients like conductance based interactions are taken into account.

In this article we present a general approach to determine the transfer function for complex models, from the Adaptive Exponential Integrate-and-Fire (AdEx) to the Hodgkin-Huxley (HH) and the Morris-Lecar (ML) models. As a result, we obtain mean-field equations for the population dynamics in AI regimes as observed in cortical regions for highly detailed models, creating a bridge between electrophysiology at the microscopic scale and the details of the famous transfer function first used by Wilson and Cowan as a sigmoid.

Finally, we test not only the ability of our mean-field models to describe spontaneous activity of the considered neuronal populations, but also their predictive power for network response to external stimuli. We show that, provided the stimuli are fairly slow, the mean field model gives good quantitative predictions.

## 2 Materials and Methods

We describe here the neuronal and network models used in this study. We also introduce mean-field equations describing population dynamics and the template to estimate the transfer function that we apply to all the neuronal models under consideration.

### 2.1 Network of spiking neurons

We consider a random directed network of *N* = 10^4^ cells, among which 80% are regular spiking (RS) excitatory (E) and 20% are fast spiking (FS) inhibitory (I) neurons. The connections between pairs of neurons are set randomly with a fixed probability (*p* = 0.05). Unless otherwise stated, the same network and synaptic constants are used for all the neuronal models (Hodgkin-Huxley, Adaptive Exponential Integrate-and-Fire and Morris-Lecar). The dynamics of each node *k* follows:

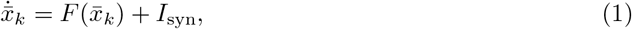

where 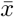 and 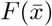 represent the neuronal state and dynamics, the latter depending on the specific model (see Section 2.2). Note the notation 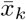 which indicates that, in general, each neuron is characterized by a vector of variables. The synaptic current impinging on the postsynaptic neurons *k*, *I*_syn_, is modeled as:

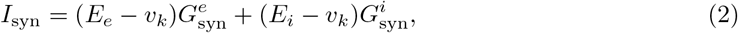

where *E*_*e*_ = 0 mV (*E*_*i*_ = −70 mV) is the excitatory (inhibitory) reversal potential and 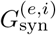 is the conductance modeled as a decaying exponential function:

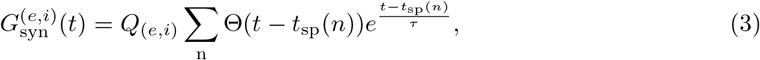

where *Q*_*e*_ (*Q*_*i*_) is the excitatory (inhibitory) quantal conductance. The variable *τ* = 5 ms is the decay timescale of excitatory and inhibitory synapses and Θ is the Heaviside step function. The summation runs over the over all the pre-synaptic spiking times *t*_sp_(*n*). For both Hodking-Huxley and Adaptive Exponential Integrate-and-Fire models we set *Q*_*e*_ = 1.5 nS and *Q*_*i*_ = 5 nS, while for Morris-Lecar model *Q*_*e*_ = 4 nS and *Q*_*i*_ = 10 nS. On top of inputs coming from other neurons in the network, each excitatory and inhibitory neuron receive an external drive in the form of a Poissonian excitatory spike train at a constant firing rate *ν*_drive_ = 4 Hz, if not stated otherwise.

### 2.2 Single neuron models

We describe here the neuronal models used in the rest of the paper, starting from the Integrate- and-Fire up to the Morris-Lecar and Hodgkin-Huxley models.

#### 2.2.1 Adaptive Exponential Integrate-and-Fire model

The dynamics of each of the AdEx neurons *i* is described by the following 2D (here 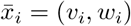) differential quations (Brette and Gerstner, 2005) :

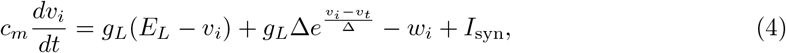

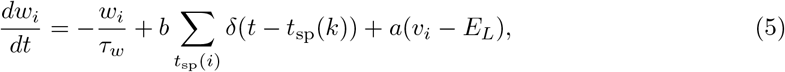

where *c*_*m*_ = 150pF is the membrane capacity, *v*_*i*_ is the voltage of neuron *i* and, whenever *v*_*i*_ > *v*_*t*_ = −50 mV at time *t*_sp_(*i*), *v*_*i*_ is reset to the resting voltage *v*_rest_ = −65 mV and fixed to this value for a refractory time *T*_refr_ = 5 ms. The leak term has a fixed conductance of *g*_*L*_ = 10 nS and the leakage reversal *E*_*L*_ = −65 mV, if not stated otherwise. The exponential term has a different strength for regular-spiking (RS) and fast-spiking (FS) cells, i.e. Δ = 2 mV (Δ = 0.5 mV) for excitatory (inhibitory) cells. The variable *w* mimicks the dynamics of spike frequency adaptation. Inhibitory neurons are modeled according to physiological insights as the FS neurons with no adaptation while the excitatory RS neurons have a lower level of excitability due to the presence of adaptation. Here we consider *b* = 60 pA, *a* = 4 nS and *τ*_*w*_ = 500 ms, if not stated otherwise.

#### 2.2.2 Hodgkin-Huxley

The dynamics of the Hodgkin-Huxley model (Hodgkin and Huxley, 1952) is given by the following 5-dimensional system of differential equations (Pospischil et al., 2008):

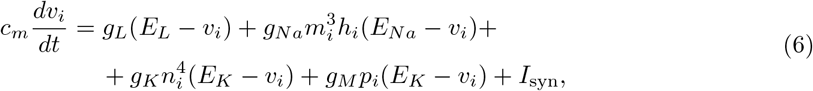

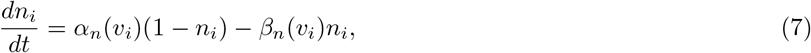

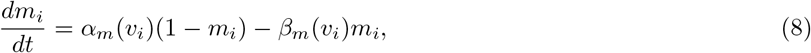

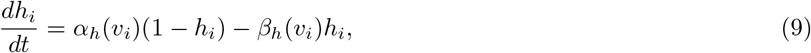

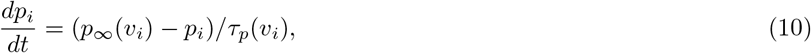

with the gating functions,

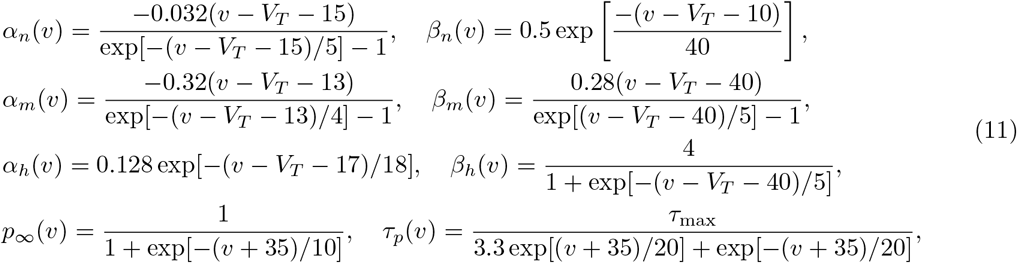

where *v*_*i*_ is the voltage and (*n*_*i*_, *m*_*i*_, *h*_*i*_, *p*_*i*_) are the corresponding gating variables of the *i*-th neuron. We set the spike emission times *t*_sp_(*k*) for this model to time steps in which the membrane potential *v* exceeded a voltage threshold of 10 mV. Unless stated otherwise, the membrane capacitance *c*_*m*_ = 200 pF/cm^2^, the maximal conductance of the leak current *g*_*L*_ = 10 mS/cm^2^, the sodium current *g*_*Na*_ = 20 mS/cm^2^, the delayed-rectifier potassium current *g*_*K*_ = 6 mS/cm^2^, the slow non-inactivating potassium current of the excitatory (RS) neurons *g*_*M*_ = 0.03 mS/cm^2^ and of the inhibitory (FS) neurons *g*_*M*_ = 0 mS/cm^2^, with corresponding reversal potentials *E*_*L*_ = −65 mV, *E*_*Na*_ = 50 mV, *E*_*K*_ = −90 mV, the spiking threshold *V*_*T*_ = −53.5 mV and *τ*_max_ = 0.4 s are the fixed parameter values in Eqs. (6)-(11).

#### 2.2.3 Morris-Lecar

The dynamics of the Morris-Lecar model (Morris and Lecar, 1981) is described by the system of offerential equations:

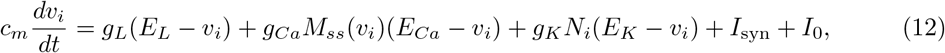

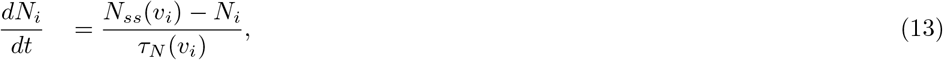

where *c*_*m*_ = 2*μ* F/cm^2^ is the membrane capacitance, *v*_*i*_ is the membrane potential in mV, *N*_*i*_ and *M*_*ss*_ are the fraction of open potassium and calcium channels, respectively. The current *I*_0_ = 0.2 nA/cm^2^ is a reference DC external current. Spike emission times are established in the same way as for the HH model. The maximal conductances for the leakage current (L), calcium (Ca) and potassium (K) were fixed to *g*_*L*_ = 20 mS/cm^2^, *g*_*Ca*_ = 80 mS/cm^2^ and *g*_*K*_ = 160 mS/cm^2^, respectively. The reversal potentials are *E*_*L*_ = −50 mV for excitatory RS neurons and *E*_*L*_ =−70 mV for inhibitory FS neurons, *E*_*Ca*_ = 120 mV and *E*_*K*_ = −84 mV. The quantities *M*_*ss*_ and *N*_*ss*_ are modeled as:

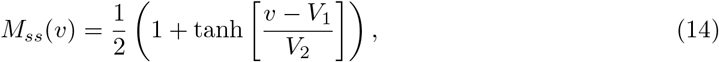

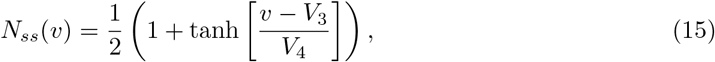

with

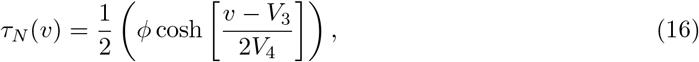

where *V*_1_ = −1.2 mV, *V*_2_ = 18 mV, *V*_3_ = 2 mV, *V*_4_ = 30 mV are tuning parameters that determine the half activating voltage and slope of the activation curves for calcium and potassium conductances. This choice of parameters is such that the ML neuron is set in a type II excitability class, i.e. its response to a DC current is discontinuous and the neuron firing rate increases very slowly with the injected current (data not shown).

### 2.3 Mean-field formalism

Mean-field theory scales the analysis of interacting point-wise neurons to their macroscopic, collective, dynamics based on the moment-statistics of the system, requiring a self-averaging hypothesis for physical quantities. We make here an additional hypothesis that the biological neural network is set to asynchronous irregular dynamical regime. The latter is chosen for its biological plausibility (Destexhe et al., 2003) as observed in awake cortical states of adult mammalian brains.

We use here the master equation formalism reported by (El Boustani and Destexhe, 2009) providing a system of ordinary differential equations that describe the evolution of the mean and variance of the firing rate of excitatory and inhibitory neurons. The central argument for this derivation is to consider the network dynamics as markovian on an infinitesimal (a time resolution T, typically 20ms) scale, as in Buice et al. (2010); Ginzburg and Sompolinsky (1994); Ohira and Cowan (1993). Moreover, such a theory is based on the assumption that neurons emit maximum one spike over the Markovian step *T*, meaning that the theory assumes relatively low firing rate of neurons, lower than 1*/T* ~ 50 Hz (El Boustani and Destexhe, 2009), as typically is the case in 123 the ynchronous irregular regimes here investigated. The differential equations read:

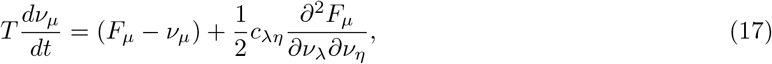

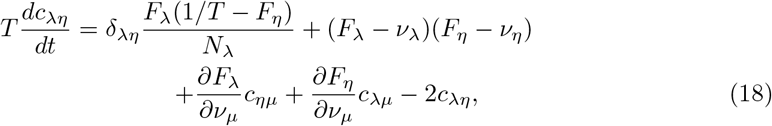

where *μ* = {*e, i*} is the population index (excitatory or inhibitory), *ν*_*μ*_ the population firing rate and *c*_*λη*_ the covariance between populations *λ* and *η*. The function *F*_*μ*={*e,i*}_ = *F*_*μ*={*e,i*}_(*ν*_*e*_, *ν*_*i*_) is the transfer function which describes the firing rate of population *μ* as a function of excitatory and inhibitory inputs (with rates *ν*_*e*_ and *ν*_*i*_). At the first order, i.e. neglecting the dynamics of the covariance terms *c*_*λη*_, this model reduces to the well known Wilson-Cowan model, with the specificity that the functions *F* need to be obtained according to the specific single neuron model under consideration. We introduce this procedure in the next section.

### 2.4 Transfer function estimate

The transfer function relates the firing rate of a neuron’s response to its presynaptic excitatory and inhibitory firing rates. The particular form of the transfer function is related to the dynamics describing neuronal activity. Deriving an analytical expression for the transfer function is a non-trivial endeavor due to the nonlinear character of the dynamics, e.g. through conductance based interactions. Therefore, we use here a semi-analytic approach to fit a family of plausible transfer functions to the data obtained by means of numerical simulations with the desired neuron types.

This method, developed first by Zerlaut et al. (2016) on data from experimental recordings, is based on the assumption that the transfer function depends only on the statistics of the subthreshold membrane voltage dynamics which is assumed to be normally distributed. These statistics are: the average membrane voltage *μ*_*V*_, its standard deviation *σ*_*V*_ and auto-correlation time *τ*_*V*_. Under these assumptions the neuronal output firing rate *F*_*ν*_ is given by the following formula:

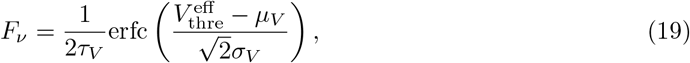

where erfc is the Gauss error function, 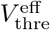 is an *effective* or *phenomenological threshold* accounting for nonlinearities in the single-neuron dynamics. Note that when dealing with extremely high spiking frequencies, e.g. in the case of Hodgkin-Huxley model close to depolarization block, a multiplicative factor *α* can be added in front of r.h.s. of Eq. (19) to permit the fitting procedure to deal with such high frequencies. In the asynchronous irregular dynamical regime, investigated in this work, neurons have relatively low firing rates (smaller than 30 Hz). Accordingly, we never use this extension (i.e. the inclusion of the factor *α*) apart from the inset of Fig. 2b where we fit the transfer function of the Hodgkin-Huxley model over a broad range of frequencies, including those close to depolarization block where the firing rate is around 500-600 Hz. For this case we used *α* = 2. In the following section we introduce how the quantities *μ*_*V*_, *σ*_*V*_, *τ*_*V*_ can be expressed as functions of the presynaptic excitatory and inhibitory firing rates *ν*_*E*_ and *ν*_*I*_.

**Table 1.** Fit Parameters AdEx Neurons (in *mV*)

| Cell Type | $P_0$ | $P_{\mu V}$ | $P_{\sigma V}$ | $P_{\tau V}$ | $P_{\mu V}^2$ | $P_{\sigma V}^2$ | $P_{\tau V}^2$ | $P_{\mu V \sigma V}$ | $P_{\mu V \tau V}$ | $P_{\sigma V \tau V}$ |
| --- | --- | --- | --- | --- | --- | --- | --- | --- | --- | --- |
| RS-Cell | -49.8 | 5.06 | -23.4 | 2.3 | -0.41 | 10.5 | -36.6 | 7.4 | 1.2 | -40.7 |
| FS-Cell | -51.5 | 4.0 | -8.35 | 0.24 | -0.50 | 1.43 | -14.7 | 4.5 | 2.8 | -15.3 |

**Table 2.** Fit Parameters Hodgkins-Huxley Neurons (in *mV*)

| Cell Type | $P_0$ | $P_{\mu V}$ | $P_{\sigma V}$ | $P_{\tau V}$ | $P_{\mu V}^2$ | $P_{\sigma V}^2$ | $P_{\tau V}^2$ | $P_{\mu V \sigma V}$ | $P_{\mu V \tau V}$ | $P_{\sigma V \tau V}$ |
| --- | --- | --- | --- | --- | --- | --- | --- | --- | --- | --- |
| RS-Cell | -48.1 | 3.2 | 10.9 | -0.32 | 0.98 | 1.1 | -1.2e-3 | -1.4 | 3.9 | -0.11 |
| FS-Cell | -51.2 | 1.8 | -6.1 | -0.86 | 1.6 | -0.70 | -11 | -0.18 | 1.2 | -1.2 |

**Table 3.**
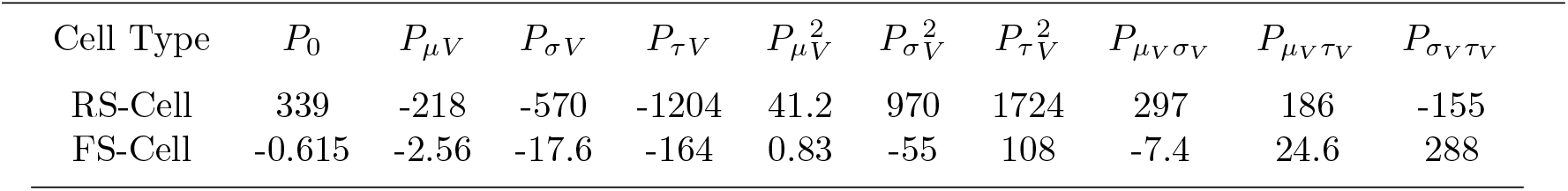
Fit Parameters Morris-Lecar Neurons (in *mV*)

| Cell Type | $P_0$ | $P_{\mu V}$ | $P_{\sigma V}$ | $P_{\tau V}$ | $P_{\mu V}^2$ | $P_{\sigma V}^2$ | $P_{\tau V}^2$ | $P_{\mu V \sigma V}$ | $P_{\mu V \tau V}$ | $P_{\sigma V \tau V}$ |
| --- | --- | --- | --- | --- | --- | --- | --- | --- | --- | --- |
| RS-Cell | 339 | -218 | -570 | -1204 | 41.2 | 970 | 1724 | 297 | 186 | -155 |
| FS-Cell | -0.615 | -2.56 | -17.6 | -164 | 0.83 | -55 | 108 | -7.4 | 24.6 | 288 |

#### 2.4.1 From input rates to sub-threshold voltage moments

We start by calculating the averages (*μ*_*Ge,Gi*_) and standard deviations (*σ*_*Ge,Gi*_) of the conductances given by Eq. 3 under the assumption that the input spike trains follow the Poissonian statistics (as is indeed the case in asynchronous irregular regimes here considered). In such case we obtain (Zerlaut and Destexhe, 2017a):

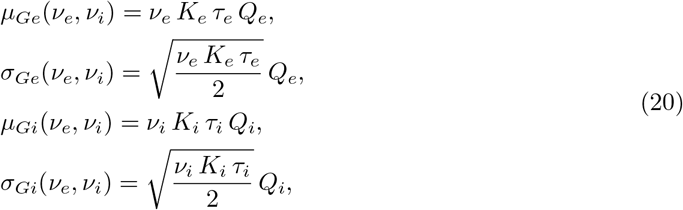

where *K*_*i,e*_ is the average input connectivity received from the excitatory or inhibitory population (in our cases typically *K*_*e*_ = 400 and *K*_*i*_ = 100) and in our model *τ*_*e*_ = *τ*_*i*_ = *τ* (see Eq. 3).

The mean conductances will control the total input of the neuron *μ*_*G*_ and therefore its effective membrane time constant 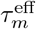:

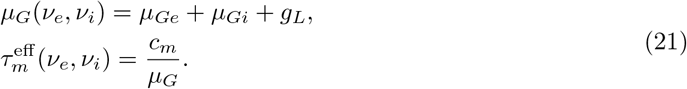

Here we make the assumption that the subthreshold moments (*μ*_*V*_, *σ*_*V*_, *τ*_*V*_) are not affected by the dynamics of the currents coming into play at the spiking time (e.g. sodium channels dynamics or the exponential term of the AdEx model). We thus consider, for all neurons, only the leakage term and the synaptic input in order to estimate subthreshold moments. Accordingly, we can write the equation for the mean subthreshold voltage:

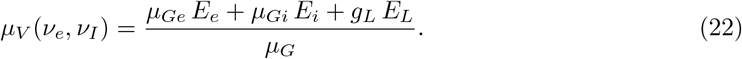

The final formulas for *σ*_*V*_ and *τ*_*V*_ follow from calculations introduced in Zerlaut et al. (2018) and they read:

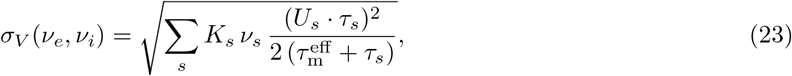

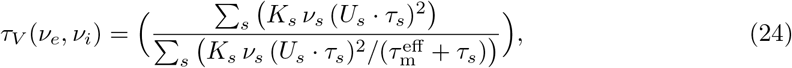

where we defined 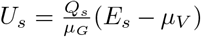 and *s* = (*e, i*). Notice that neglecting all the spiking currents becomes a poorer assumption as the neuron activity increases. Nevertheless, we consider here AI dynamics where neurons have typically low firing rates (of the order few Hz). Moreover, as we show in the following sections, the fitting procedure will account for discrepancies in the actual evaluation of voltage moments by permitting an accurate prediction of neuron output firing rate.

#### 2.4.2 From sub-threshold voltage moments to the output firing rate

The quantities *μ*_*V*_, *σ*_*V*_ and *τ*_*V*_, obtained in the previous section, can now be plugged into Eq. 19 when an additional relation is taken into account. This relation follows from theoretical and experimental considerations (Zerlaut et al., 2016) showing that the voltage effective threshold 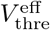 can be expressed as a function of (*μ*_*V*_, *σ*_*V*_, *τ*_*V*_). In Zerlaut et al. (2016) the phenomenological threshold was taken as a second order polynomial in the following form:

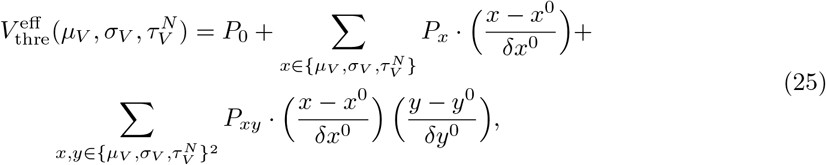

where we introduced the quantity 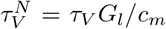. We evaluated {*P*} through a fit according to simulations on single neurons activity setting first 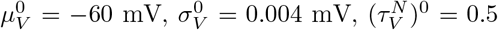, 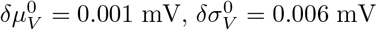 and 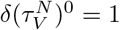. By the fitting procedure we find the values of the P parameters for the three neuronal models considered here (additionally for each model we consider two neuronal types: RS and FS) and we report the results in Table 3. In the first part of the Results section we describe the goodness of this procedure for the three considered neuronal models.

## 3 Results

We present here the results of a comparison between mean-field predictions and direct simulations. We first test the technique to estimate the transfer function of single cells in AdEx, Hodgkin-Huxley and Morris-Lecar models and then compare theoretical predictions of the mean-field to numerical simulation of sufficiently large networks of neurons.

### 3.1 Transfer function for integrate-and-fire models

The transfer function of a simple AdEx neuron can be straightforwardly estimated by numerical simulations. As we report in Fig. 1 its shape is very similar to a sigmoidal function (as in the seminal paper by Wilson and Cowan) but its specific parameters follow from a complex combination of microscopic information, e.g. neurons resting potential. See the black and blue dots in Fig. 1 for different values of the leakage reversal potential *E*_*L*_. Two main spiking modes can be distinguished in the neuronal dynamics. One, characterized by low output firing rate, where spikes are strongly driven by the membrane voltage fluctuations, namely fluctuation driven mode (see the bottom inset in Fig. 1). The second mode is characterized by a highly deterministic and regular firing observed at very high output firing rates (larger than 40 − 50 Hz, top inset).

**Figure 1.**
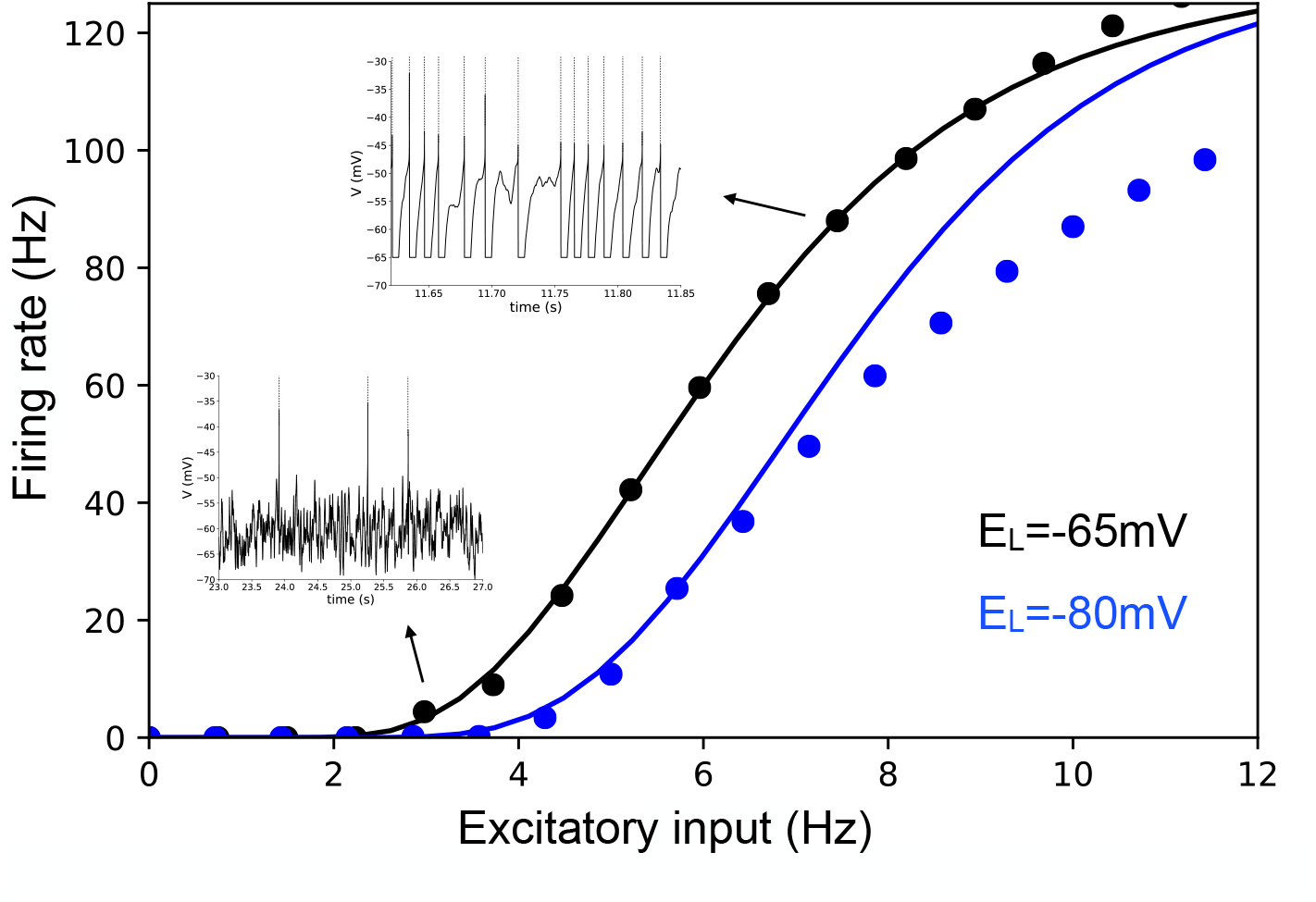
Transfer function for an Exponential Integrate-and-Fire model. Dots indicate the results of the numerical simulation of the Exponential Integrate-and-Fire model (FS cell, see Materials and Methods). The continuous line illustrates the results based on the semi-analytic fitting. The inhibitory Poissonian spike train used here has a fixed rate, *r*_*I*_ = 8 Hz, while we show neuron average output Firing rate as the function of the Poissonian excitatory input spike train of rate *r*_*E*_. In the insets we show two exemplary voltage time traces corresponding to high (top inset) and low (bottom inset) firing rate. Colors stand for different values of the leakage reversal potential as indicated in the bottom-right corner of the figure.

By employing the semi-analytic approach to predict the transfer function we observe a very good agreement with direct simulations (see continuous lines in Fig. 1, showing predictions based on this approach) as it has been shown by El Boustani and Destexhe (2009); Zerlaut et al. (2016). The agreement remains very good for relatively low neuronal activity (up to 50 Hz). This is a direct consequence of the semi-analytic approach that assumes that neurons fire in an irregular manner (as observed in cortical dynamics) strongly driven by fluctuations around the mean membrane voltage. In this work we only consider Asynchronous Irregular population dynamics for which the activity of neurons is low, irregular and strongly fluctuation driven.

### 3.2 Transfer function for complex models

We report here the application of the techniques described in the Materials and Methods section to evaluating the transfer function of more complex neuronal models. To this end we consider the well-known Hodgkin-Huxley (HH) model and the Morris-Lecar (ML) models (see Materials and Methods). These models permit to describe the details of Sodium and Potassium channels dynamics neglected in the simpler Integrate-and-Fire model and reproduce time evolution of the action potential. The semi-analytic approach to fit the numerical transfer function can be applied exactly in the same way as for AdEx models (as discussed in the Materials and Methods section).

We consider two kinds of neurons in agreement with neurophysiolgical information about cortical cells: excitatory neurons modeled as RS cells with a lower gain of the transfer function and inhibitory neurons modeled as FS cells with a higher gain. A different gain of the transfer function can be obtained by changing the excitability of the cells through their resting potential, or increasing the adaptation strength (see Materials and Methods for details).

By comparing the theoretical prediction with numerical simulation we observe that, for the three models considered here, the transfer function is correctly estimated both for inhibitory neurons (FS cells) and excitatory neurons (RS cells). This result shows that, even by considering a much more complicated model than AdEx it is possible to have access to a semi-analytic form of its transfer function and, importantly, to modify neurons excitability in the way allowing to obtain a similar transfer function (of excitatory RS and inhibitory FS cells) between different models.

Notice that the ML model shows a decrease of firing rate at frequencies higher than 8 Hz (i.e. no voltage oscillations and thus no firing activity), as reported previously for this model by Kim and Nykamp (2017). This is a consequence of the depolarization block (DB) observed at high input frequencies (i.e. high average external current). Accordingly, we obtain a bell shaped transfer function, well predicted by our semi-analytical formalism. In previous studies this effect was taken into account in the context of Wilson-Cowan equations by using a Gaussian transfer function, instead of a sigmoidal Meijer et al. (2015), permitting to study the effect of the depolarization block in focal seizures at the population scale. In our model this shape, resembling a Gaussian, follows directly from Morris-Lecar equations, through the semi-analytical fitting. More specifically, in Meijer et al. (2015) the DB was studied in the Hodgkin-Huxley model. Indeed, also the HH model shows a DB but, at variance with the Morris-Lecar case, it appears in our parameters setup at very high firing rates, around 600-700 Hz (see the inset of Fig. 2b). In our simulations we do not consider this dynamical regime, being far from the dynamics typical of neurons in asynchronous irregular regimes. Moreover, as described in Fig.1, the semi-analytic fitting procedure works well for low firing rates and discrepancies appear at high rates, also in the case of the simpler AdEx model. Nevertheless, we report here that, when performing the fitting over a wide range of input and output rates (see Methods), it is possible to obtain an overall good fit of the bell-shaped transfer function (see the inset of Fig. 2b).

**Figure 2.**
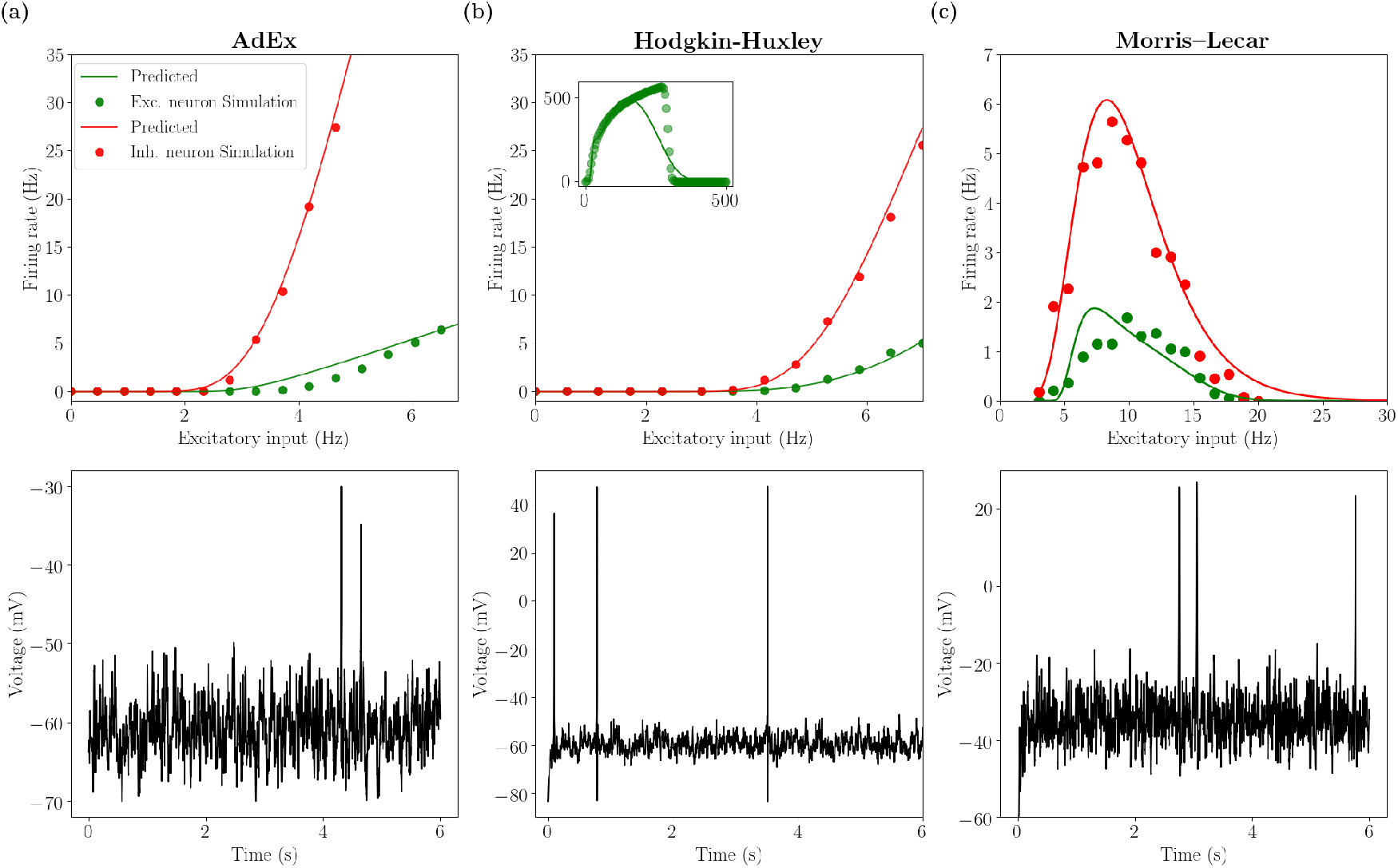
Transfer function for RS and FS cells: AdEx, HH and ML. We report the output firing rate for excitatory RS (green) and inhibitory FS (red) cells obtained from numerical simulation (dots) and from the semi-analytic approach for the transfer function (continuous line). The inhibitory Poissonian spike train has a fixed rate, *r*_*I*_ = 8 Hz. The bottom panels shows the time traces of the membrane voltage of an RS cell for an excitatory input equal to 4 Hz. Left column is obtained for the AdEx model, middle column for the HH model and right column for the ML model (see Materials and Methods). In the inset of panel (b) we report the transfer function for the RS cell estimated over very large values of input rates. In this case a separate fit by considering a broad input frequency range has been performed (see Materials and Methods).

Beyond the methodological point, our results show that even if the details of the mechanisms that generate a specific transfer function are very different, it is possible to adjust neurons parameters (e.g. excitability) in a way allowing to obtain similar transfer functions (at least in the region before entering a depolarization block). As a consequence, according to the mean-field theory, where what matters for the population dynamics is only the transfer function, we expect different models to have a comparable emergent dynamics at the population (collective) scale.

### 3.3 Asynchronous irregular dynamics and mean-field predictions

In this section we compare the mean-field predictions of the emergent dynamics of networks of AdEx, HH and ML neurons, in the setup described in the previous section. In particular, we simulate a sparse network of RS and FS cells (see Fig. 2) coupled through conductance based interactions (see Materials and Methods for details). By looking at Fig. 2 we observe that, before reaching the DB, all three models have similar transfer functions, with FS neurons having a higher gain with respect to RS neurons, approximately of factor 3–4. As a result, we expect the population activity in the three models to fall within a similar dynamical regime, as a natural consequence of the mean-field’s sole dependence on transfer functions, previously stated.

Indeed, by looking at Fig. 3 we observe that in the different networks the dynamics stabilizes on an asynchronous regime. In all cases, this regime is characterized by irregular microscopic dynamics (neuron’s spiking statistics are Poissonian, data not shown) and represents the typical spiking patterns recorded during awake states in cortical regions (the autocorrelation function of population rate decreasing to zero in the time scale of tens of milliseconds). Moreover, as expected, inhibitory FS cells fire at a higher frequency with respect to RS cells. Through the mean-field model it is possible to measure both the average population rate and its covariance (second order mean-field, see Materials and Methods). As reported in Fig. 3 we show that the mean-field model gives a good quantitative prediction of both quantities when they are compared to the histogram obtained by sampling the population rate in the network simulation. The higher discrepancy we observe for the complex neuronal models (e.g. HH and ML case) is related to a higher mismatch of the transfer function linked to the higher complexity of the model.

**Figure 3.**
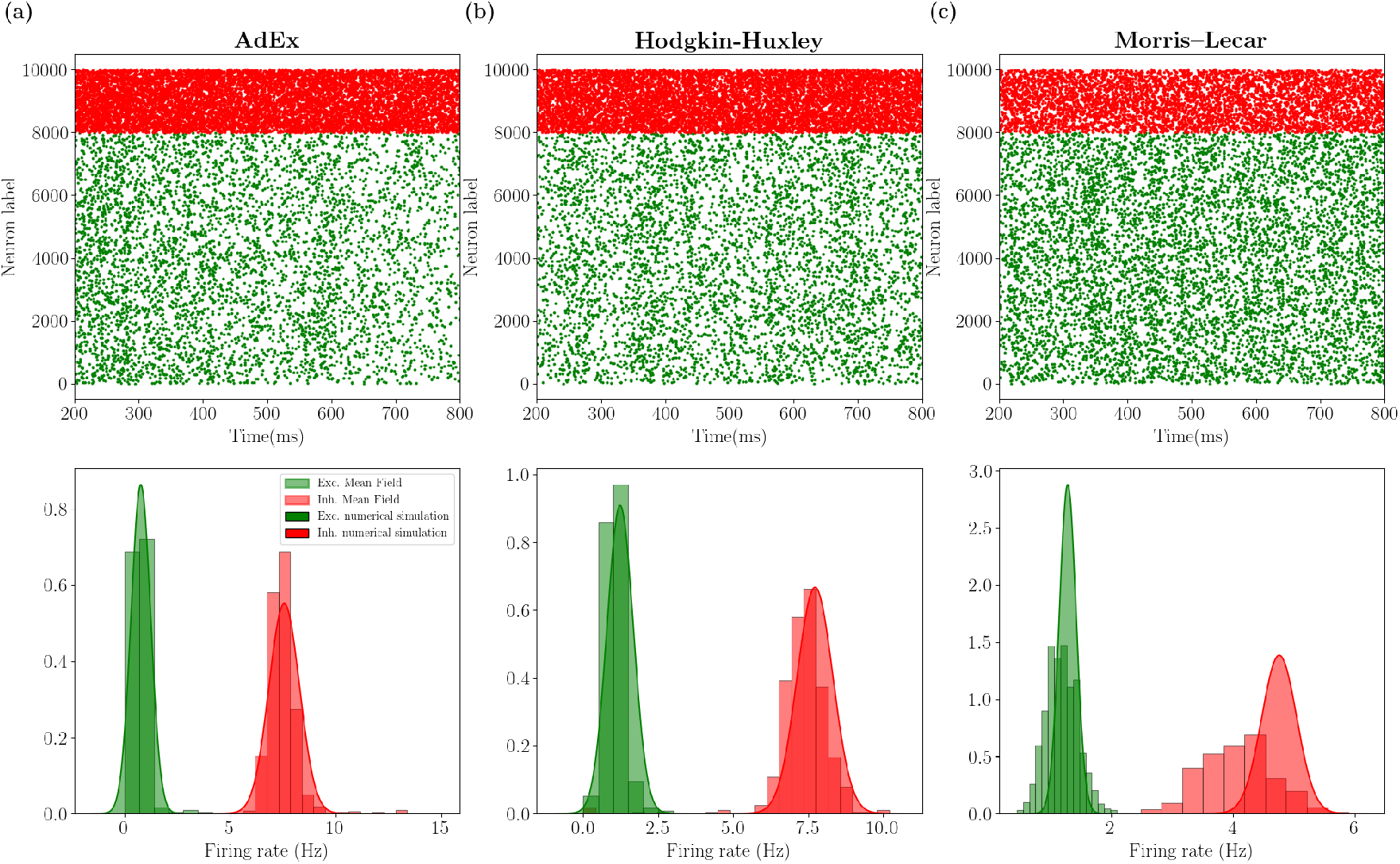
Mean-field predictions and spontaneous activity: AdEx, HH and ML. Top panels show raster plots for excitatory (green dots) and inhibitory (red dots) neurons, i.e. the spiking times for each neuron. Bottom panels shows histograms (obtained on a time length *T*_*w*_ = 10 s) of population firing rated for excitatory (green) and inhibitory (red) populations. The Gaussian distribution has been plotted from mean-field predictions giving access to average firing rate and its variance. The left column (a) is obtained for the AdEx model, the middle column (b) for the HH model and the right column (c) for the ML model (see Materials and Methods).

### 3.4 Network response to external stimuli

In order to complete the comparison between the mean-field model and the network dynamics, we study the response of the system to external stimuli. In particular, we consider an incoming Poissonian train of spikes characterized by time-varying frequency and targeting both excitatory and inhibitory cells according to the following equation:

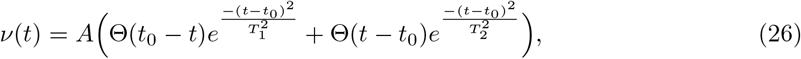

where Θ is the Heaviside function, and *T*_1_ and *T*_2_ are the rise and decay time constants, respectively. In Fig. 4 we report the comparison between the mean-field prediction and the network dynamics.

**Figure 4.**
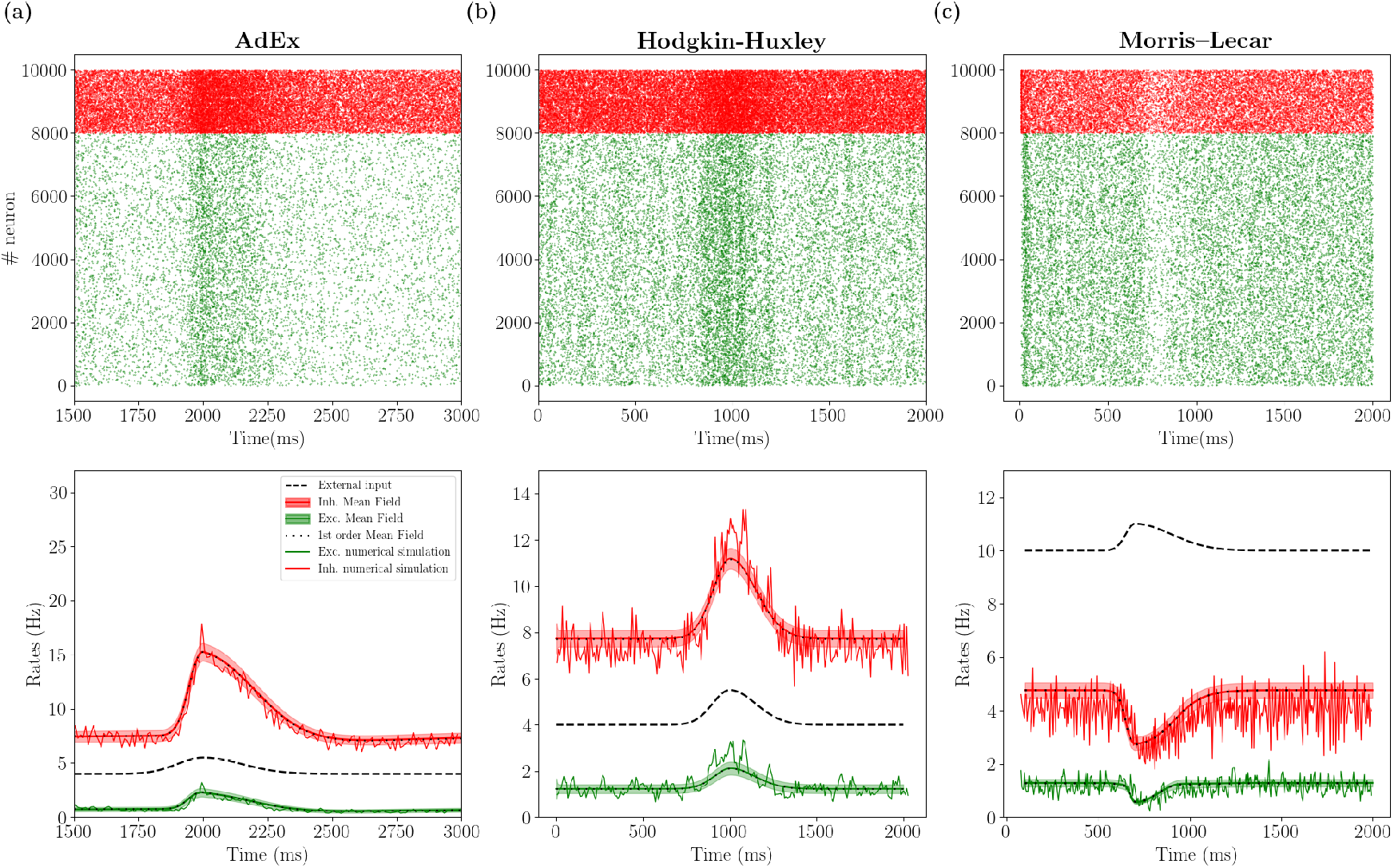
Population response to external stimuli: AdEx, HH and ML. Top panels show the raster plot for excitatory (green dots) and inhibitory (red dots) neurons in response to an external excitatory stimulus (black dashed line in lower panels). Lower panels show the corresponding population rate (noisy line) together with the mean and standard deviation over time predicted by the the second order mean field model (red for inhibition and green for excitation). Superimposed the result obtained for the mean-field at the first order (black dots), which are almost coincident with results at the second order. Left column is obtained for the AdEx model, middle column for the HH model and right column for the ML model. Parameters are the same as in Fig. 3 and the external input (see Eq. (26)) has parameters *A* = 2 Hz, *T*_1_ = 100 ms, *T*_2_ = 150 ms for AdEx and HH and *A* = 2 Hz, *T*_1_ = 100 ms, *T*_2_ = 150 ms for ML, with *t*_0_ = 2 s.

By looking first at the AdEx and HH models, we observe that both mean-field models under investigation compare favorably with their corresponding network dynamics. We also verified, as it has been shown in (di Volo et al., 2019), that the faster the input dynamics is, the worse the agreement becomes. Indeed, for the Markovian hypothesis to hold, we need the time scale *T* to be much larger than the autocorrelation time in the spontaneous activity *T* ~ *τ*_*m*_ ~ 10 ms.

Considering now the case of the ML model, we observe that by looking at Fig. 2 a relatively strong input would bring single neurons to a depolarization block, which appears at relatively low activity levels. According to this difference with respect to AdEx and HH models, we would expect the population dynamics to show different properties in response to external perturbations. Indeed, as reported in Fig. 4c, the response to an external stimulus is very different from the one observed in the HH and AdEx models. In fact, in this case the excitatory stimuli turns out to inhibit both population activities. This anti-correlation between population input and output is well captured in its time course also by the mean-field model. This result shows that also for a more complex and highly non-linear setup the mean-field model is capable of predicting the ongoing activity and the time course of the response of a network of neurons operating in the asynchronous irregular dynamical regime.

Finally, we compare the results of the first and second order mean field on average population rates. In Fig. 4 we superimpose the continuous green (red) line for excitatory (inhibitory) rate obtained with the second order mean file with the results obtained with the first order (black dots). We observe that the two quantities almost overlap (the difference is too small to be appreciated at this scale). Nevertheless, it is worth noticing that the second order mean field permits to obtain non-trivial information on the population dynamics and its fluctuations in time, with good quantitative predictions of the covariance of population rates (see the histograms in Fig. 3 and shadows in Fig. 4)).

## 4 Discussion

In this manuscript, we have reviewed a formalism to derive mean-field models from networks of spiking neurons and we have applied it to different complex neuronal models. The key to derive such “biologically realistic” mean-field models is to be able to obtain the transfer function of complex neuronal models. The approach we followed used a mean-field formalism based on a Master Equation and which is applicable to every neuron, provided the transfer function is known (El Boustani and Destexhe, 2009). More recently, we have shown that the usual mathematical form of the transfer function, known analytically for the Integrate-and-Fire model, can capture more complex neuronal models (Zerlaut et al., 2018, 2016). This gave rise to a “semi-analytic” approach, where the transfer function is parameterized and fit numerically to the neuron model, while the mean-field remains analytic as only the parameters are obtained from the fitting. This approach was applied to the AdEx model (di Volo et al., 2019; Zerlaut et al., 2018), and we extend it here to more complex models, namely the Morris-Lecar and the Hodgkin-Huxley models.

It is important to note that we limited here to “simple” firing patterns, i.e. neurons fire tonically in response to an external stimulus. In this setup the transfer function is well defined as the neuron’s firing rate defines completely the spiking pattern. In cases where neurons exhibit different kind of activity, e.g. bursting, a different approach needs to be employed (see (Ostojic and Brunel, 2011)). Nevertheless, in the context of tonic neuronal activity the method is shown to be able to capture the response function of highly realistic models. We have studied here the predictions of the considered mean-field models on networks dynamics of excitatory RS and inhibitory FS cell populations during asynchronous irregular regimes, as observed in awake cortical activity. The results positively compare in the case of Morris-Lecar and Hodgkin-Huxley models for both the average and the variance of network population activity.

The good predictions at the population levels in the framework of the asynchronous irregular regimes is strongly dependent on the goodness of the fitting procedure for single neurons transfer functions. Even if such procedure works very well for neurons working in a low rate regime, whenever the firing rate becomes very high (higher than 100 Hz) the quantitative agreement gets worse. A more refined technique for the evaluation of the transfer function in different states (low and high rates activity) is an important topic for future research (recent work has addressed this issue in AdEx model (Capone et al., 2019a)). A step forward in this direction can be important when dealing with neurons entering depolarization block at high firing rates, a mechanism playing an important role in focal seizures (Meijer et al., 2015) or in dopaminergic neurons under normal condition or under drugs assumption (Di Volo et al., 2019; Dovzhenok and Kuznetsov, 2012). Both in Morris Lecar and Hodgkin-Huxley model, the semi-analytic fitting is found to give quite good predictions on the presence of depolarization block, especially in the Morris Lecar case as this setup does not consider very high spiking frequencies. Even if work remains to be done in order to extend this framework to obtain a more reliable quantitative prediction on the depolarization block at very high frequencies, these preliminary results indicate the possibility to use these mean field techniques to connect the physiology at the cellular scale with pathological states at the population level, as the case of focal seizures.

We also reported that, in the framework of the Asynchronous regimes here considered, corrections to first order mean field due to second order terms (see Eq. (17)-(18)) were relatively small but gave a good quantitative indication on the covariance of population rates (see histograms in Fig. 2). Nevertheless, in the case of dynamical regimes with higher neuronal correlation with respect to the ones here considered, we expect the second order mean field (taking explicitly into account the dynamics of covariances), to play an important role in the prediction of population average collective dynamics.

Let us point out that the goodness of the mean-field prediction depends also on the emergent dynamics of the network, i.e. in a highly synchronous dynamical regime the Markovian hypothesis fails and the mean-field model cannot give accurate predictions. Nevertheless, even if light synchronization is considered, e.g. during slow waves sleep, the mean-field models has been shown to correctly predict such collective oscillations (di Volo et al., 2019). In this case it is however necessary to consider a mean-field model that includes the slow dynamics of spike frequency adaptation or that of the *I*_*M*_ current in the case of Hodgkin-Huxley model. The possibility to include a conductance based adaptation to this formalism, e.g. by considering the slow dynamics of *I*_*M*_ current, is a stimulating perspective for future works and will permit to obtain mean-field models for realistic neuronal models beyond the asynchronous irregular regime.

Moreover, beyond the input-output transfer function used here, a more complex transfer function has been used in order to take into account other features of neuron response, e.g. response in frequency (Ostojic and Brunel, 2011). The addition of variables to account for a richer spiking pattern is an interesting direction, in case one is interested in modeling brain regions characterized by non-tonic firing of neurons (e.g. bursting cells in the thalamus). The general framework presented here could be extended in this direction, as it has been done to account for spike frequency adaptation yielding slow oscillations at the population scale.

Another possible extension is to apply the same formalism to complex neuronal models that include dendrites. A first attempt has been made in this direction (Zerlaut and Destexhe, 2017b) by considering simple “ball and stick” neuron models, where some analytic approximation is possible. In principle, it should be possible to apply this approach to models based on morphologically-reconstructed neurons, and calculate the transfer function of such models (work in progress). This will lead to mean-field models based on morphologically realistic neuronal models. However, the presence of dendritic voltage-dependent currents complicates this approach, and should be integrated in the formalism. This constitutes an exciting future development of our approach.

Finally, the positive results obtained here for complex models, by showing the generality of our approach, motivate the future step of the application of this technique directly to experimental data. To this end, neurons must be recorded intracellularly in the absence of network activity (as typically *in vitro*), and many combinations of excitatory and inhibitory inputs must be injected as conductances (using the dynamic-clamp technique). The first attempt of this sort was realized on the Layer 5 neurons from mouse primary visual cortex (Zerlaut et al., 2016), where the transfer function could be reconstructed for a few dozen neurons. The same dynamic-clamp experiments should be done in the future to characterize the transfer function of inhibitory interneurons. Based on such experiments, it will be possible to obtain a mean-field model based on the properties of real neurons. Such a model will evidently be more realistic than the models we have presented here, which must be considered as a first step towards a quantitative population modeling of cerebral cortex and other brain regions.

## 5 Acknowledgments

This research resulted from student projects during the Spring School of the European Institute of Theoretical Neuroscience (www.eitn.org), of which the participating students signed as co-first authors here. Research supported by the CNRS and the European Union (Hunan Brain Project H2020-720270 and H2020-785907). M.J. acknowledges support from the European Research Council under the European Union’s Seventh Framework Programme (FP/2007-2013)/ERC Grant Agreement No. 616268 F-TRACT and the European Unions Horizon 2020 Framework Programme for Research and Innovation under Specific Grant Agreement No. 785907 (Human Brain Project SGA2).

